# UFMylation suppresses Type IFN signaling during *Mycobacterium tuberculosis* infection of human macrophages

**DOI:** 10.1101/2024.08.07.607094

**Authors:** Nicholas E. Garelis, Rutger D. Luteijn, David H. Raulet, Jeffery S. Cox

## Abstract

Type I Interferons (IFN-I) promote host defense against a wide range of viral infections, but inhibit control of several bacterial pathogens, including *Mycobacterium tuberculosis* (Mtb). Given the significance of IFN-I signaling in determining Mtb infection outcomes, we sought to uncover new molecular mechanisms that regulate IFN-I during mycobacterial infection, specifically focusing on how IFN-I signaling is regulated at the earliest stages of mycobacterial infection as modeled by macrophage infection. To comprehensively identify these host genes, we performed a reporter-based, genome- wide CRISPR-interference screen in human macrophages infected with *Mycobacterium marinum*, a close relative of Mtb. Our screen detected 2035 significantly enriched genes (p<0.05), which included many known regulators of IFN-I, but also many unexpected genes as potentially novel regulators of IFN-I signaling. One of these unexpected genes was UFL1, an E3-like ligase that catalyzes conjugation of the ubiquitin-like protein, UFM1, in a process termed UFMylation. UFL1-deficiency during Mtb macrophage infection resulted in increased expression of IFN-β, interferon-stimulated genes, and other pro-inflammatory genes, such as TNF and IL-6, at both the transcript and protein level. Depletion of other UFMylation components phenocopied UFL1-deficiency, suggesting that UFMylation activity is required for IFN-I repression. Full transcriptional profiling revealed a broad increase in the inflammatory response of UFL1-deficient cells during Mtb infection, including both protective inflammatory cytokines and potentially detrimental interferon-stimulated genes. Our results suggest a role for UFMylation in suppressing IFN-I signaling and inflammatory responses during the earliest stages of Mtb infection.

**IMPORTANCE:** *Mycobacterium tuberculosis* was estimated to have caused over 1 million deaths in 2025 – the most deaths globally by a single bacterial pathogen. Past studies in mice and humans have shown that some host immune responses, such as IFN-I signaling, can enhance susceptibility to *M. tuberculosis*. We report the results of our genome-wide CRISPR-interference screen to determine regulators of IFN-I signaling during the earliest stages of mycobacterial infection. This screen identified new regulators of IFN-I signaling, including the UFMylation pathway, which appears to play an unexpected role in suppressing both IFN-I signaling and a broader inflammatory response. These findings may be relevant for the development of therapeutics and prophylactics for tuberculosis that impact IFN-I signaling – a known determinant of tuberculosis susceptibility.

## INTRODUCTION

*Mycobacterium tuberculosis* is responsible for the most deaths globally by a single bacterial pathogen, and its pathogenesis is driven by both bacterial factors as well as deleterious host responses (1, 2). In particular, IFN-I, including IFN-β and several IFN- αs, comprise a family of host immune signaling molecules that are protective in the context of many viral infections and certain bacterial infections, but potently inhibit some antibacterial host responses and exacerbate infection for some pathogenic bacteria, including Mtb (3–16).

The detrimental role for IFN-I during Mtb infection is supported by a wide-range of human studies: interferon signaling correlates with active tuberculosis disease, and transcriptional signatures for IFN-I signaling preceded transition to active tuberculosis disease by 18 months (17, 18). Alleles of the IFN-I receptor subunit IFNAR1 that reduce IFN-I signaling also correlated with lower tuberculosis lung pathology (11). Additionally, when hepatitis patients were treated with IFN-I therapy, there were occurrences of reactivation of Mtb (19). Thus, past studies provide a compelling case for a detrimental role for IFN-I during human Mtb infection.

A detrimental role for IFN-I during Mtb infection has also been recapitulated in mouse models: Mice lacking IFNAR1 are resistant to tuberculosis, with particularly strong effects observed in a highly-Mtb-susceptible mouse model (7, 9). The precise mechanism by which IFN-I enhance Mtb susceptibility remains an active area of investigation, but, to date, several effects of IFN-I have been implicated as potential mechanisms, including the impairment of signaling by other cytokines – particularly IFN-γ, IL-1, and TNF – and the promotion of cell death (4, 5, 9, 15, 16, 20–22). In total, findings from both human and mouse studies demonstrate that IFN-I hinder control of Mtb infection.

How Mtb induces IFN-I throughout the course of host infection remains an active area of investigation, and may involve different pathways activated at different stages of infection. In the earliest stages of pulmonary Mtb infection, one of the first cell-types infected by Mtb is macrophages, which are known to produce IFN-I during early Mtb infection (4, 23). *Ex vivo* macrophage studies have demonstrated that Mtb has the property of inducing IFN-I via a bacterial virulence factor ESX-1, a type VII secretion system (7). ESX-1 activity appears to trigger the host cGAS-STING-IRF3 pathway, in which cGAS senses DNA in the host cytosol and initiates IFN-I production – although the precise mechanism by which ESX-1 activates cGAS remains poorly understood (7, 24, 25). Importantly, both ESX-1 and the cGAS-STING pathway are required for induction of IFN-I at the earliest stages of macrophage infection (7, 24). Additionally, ESX-1 is critical for full Mtb virulence, and an intact ESX-1 locus is one of the key differences between virulent Mtb and the attenuated vaccine strain, BCG (26, 27). What makes ESX-1 required for full Mtb virulence remains poorly understood, but its ability to trigger IFN-I production, a host-detrimental immune response, raises the possibility that induction of IFN-I is a primary function of ESX-1. Overall, while there are many complexities surrounding Mtb’s elicitation of IFN-I throughout the course of infection, macrophage studies suggest stimulation of cGAS-STING-IRF3 via ESX-1 elicits IFN-I during the earliest stages of Mtb infection.

After the earliest stages of Mtb infection, other host pathways likely contribute significantly to IFN-I production during the course of Mtb infection: TLR7/9, which sense nucleic acids; the RIG-I/MAVS pathway that recognizes RNA; the Nod receptors that recognize cell wall components; and TLR2 that recognizes lipids and glycoproteins from the mycobacterial cell envelope (20). Importantly, these pathways, along with the cGAS-STING-IRF3 pathway, converge on the synthesis and secretion of IFN-I which further amplify the response by signaling through the IFN-I receptor, thereby triggering additional production of IFN-I. While many regulators of these pathways and cGAS-STING signaling have been discovered – and, in several cases, have been demonstrated to be important determinants for Mtb control – our knowledge is incomplete and it seems likely that there are undiscovered pathways that regulate production of IFN-I (20, 24, 28).

UFMylation is a ubiquitin-like post-translational modification whereby the ubiquitin-like UFM1 protein is covalently attached to target substrates via an E1 (UBA5), E2 (UFC1), and E3 ligase (UFL1) system (29). There are also additional components for UFMylation, such as DDRGK1, which functions to recruit the UFMylation ligase, UFL1, to the endoplasmic reticulum (ER), where both DDRGK1 and UFL1 are reported to localize (30). UFMylation remains incompletely understood, but it has been recently implicated in maintaining ER homeostasis and triggering autophagy of the ER to resolve stalled ER-bound ribosomes (29, 30).

There are a growing number of reports suggesting a connection between UFMylation and IFN-I in the context of viral infection, although UFL1 repression of IFN-β by UFMylation during bacterial infection has not been previously reported to our knowledge (31). UFMylation’s link to ER homeostasis provides one potential mechanism for this connection, as ER function has been linked to immune responses via the unfolded protein response (UPR) (32). The UPR is activated by the accumulation of unfolded proteins within the ER lumen and it is mediated primarily through three transcription factors, XBP1, ATF6f, and ATF4, which have a range of effects, including the upregulation of chaperones and the promotion of degradation of misfolded proteins, but also the modulation of immune responses (33).

Due to consistent findings that IFN-I signaling is an important determinant of Mtb infection outcomes, we sought to uncover additional factors that regulate IFN-I – focusing on the earliest stages of mycobacterial infection as modeled by macrophage infection. Here we report the results of a genome-wide CRISPR-interference screen to identify new regulators of IFN-β during mycobacterial infection of human macrophages. This screen identified several new factors, including several components of the UFMylation pathway. We found that UFMylation-deficiency led to hyper-induction of IFN-β, interferon-stimulated genes (ISGs), and several inflammatory cytokines, in response to Mtb infection. Our findings suggest that UFMylation, a pathway for ER homeostasis, plays an unexpected role in suppressing IFN-I signaling and a broader inflammatory response in the earliest stages of Mtb macrophage infection.

## RESULTS

To methodically uncover host genes that regulate IFN-β during mycobacterial infection, we conducted a genome-wide CRISPRi screen using a previously developed THP-1 IFN-β-reporter cell line (34). This human monocyte cell line can be differentiated into macrophages and contains an ISRE-IFN-β reporter gene consisting of an interferon-beta promoter and multiple interferon-stimulated response element (ISRE) sequences fused to fluorescent tdTomato (Fig. 1a). For initial experiments, *Mycobacterium marinum* (Mm), a common model for Mtb, was used as Mm recapitulates key aspects of IFN-I signaling during Mtb infection, such as dependence on ESX-1 and STING, for robust induction of IFN-I (35–37). A multiplicity of infection (MOI) of 5 with Mm robustly induced the reporter 24 hours post infection (hpi) (Fig. 1b). CRISPRi-mediated silencing of STING and infection with Δ*esx1* (ΔRD1) mycobacteria reduced reporter signal to near basal levels, demonstrating the system faithfully recapitulated key properties of IFN-β induction by Mtb (Fig. 1b).

**Fig. 1.**
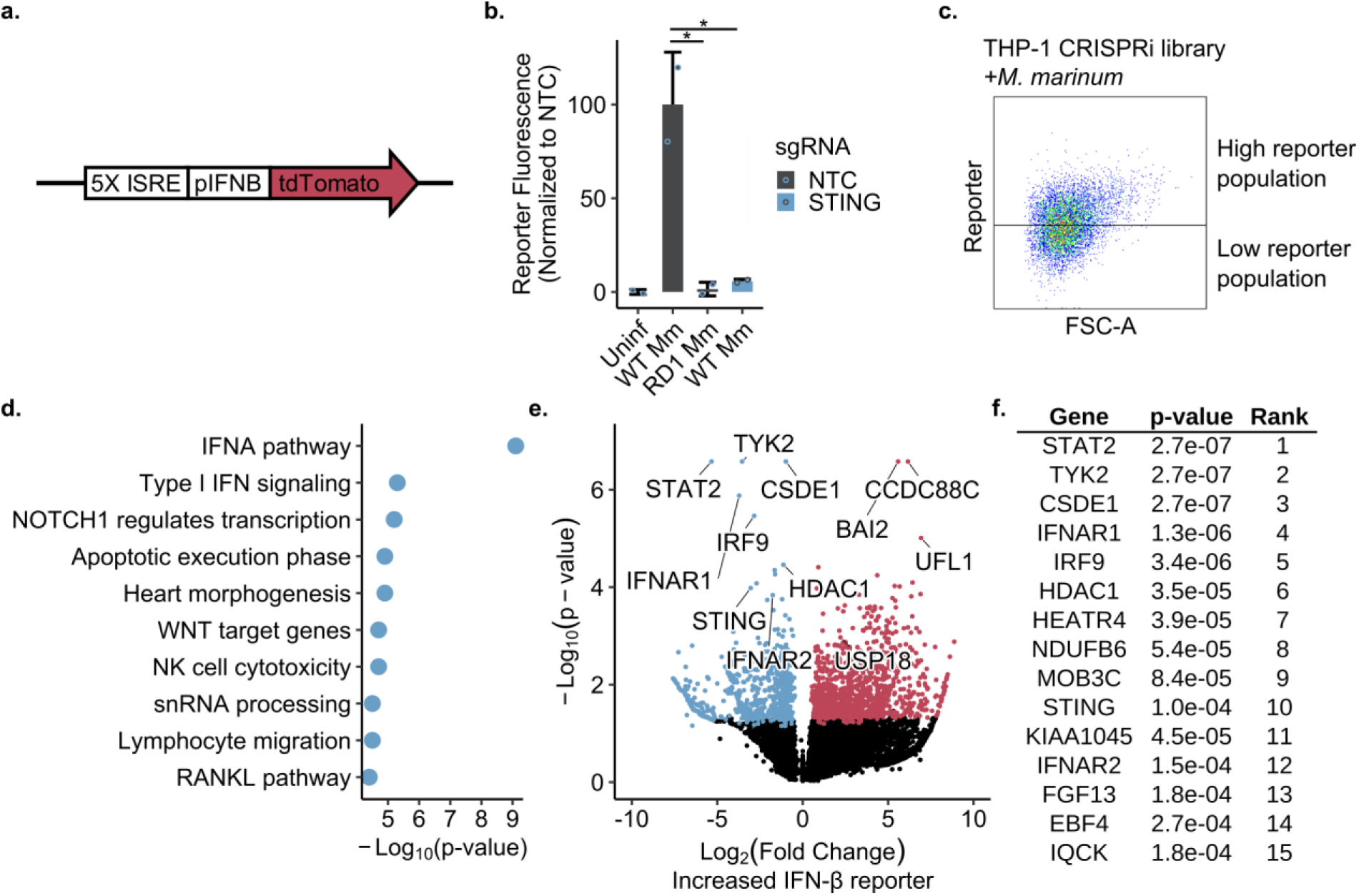
CRISPR interference screen for regulators of IFN-β during mycobacterial infection. (**a**) Schematic of the type I interferon genetic reporter with tdTomato expression driven by the IFN-β promoter and interferon-stimulatory response elements (ISREs). (**b**) THP-1 macrophages with the indicated sgRNA were infected with *M. marinum* of the indicated genetic background. At 24 hpi, reporter expression was measured by flow cytometry. Results are representative of at least two independent biological experiments. Error bars represent SD with replicates n = 2. NTC is Non- Targeting Control. (**c**). A CRISPRi library of sgRNA targeting approximately 20,000 genes at 5 sgRNA/gene was transduced into THP-1 reporter cells and cells were infected with *M. marinum*. Cells were sorted into the top 50% (high reporter population) and bottom 50% (low reporter population) as measured by reporter fluorescence for gDNA extraction. Gate in graphic is a reconstruction for illustrative purposes. (**d**). Pathway analysis of CRISPRi results, testing for pathway enrichment in the top 250 genes for the reporter low population as ranked by Robust Rank Aggregation (RRA) with MAGeCK. Some pathway names were shortened/simplified for brevity and some redundant pathways were removed or combined, keeping only the most statistically strong representative pathway. (**e**). Volcano plot showing results from the CRISPRi screen using p-values and fold-change from the MAGeCK analysis package, using results from 3 independent experiments. Red and blue points indicate MAGeCK’s rank <1000 genes. (**f**) Top genes enriched in the low-reporter population as ranked by MAGeCK’s Robust Rank Aggregation. *p<0.05, **p<0.01, ***p<0.001, ****p<0.0001 by unpaired t-test.

Next, we conducted a genome-wide CRISPR-interference screen to identify new regulators of IFN-β during infection. We generated a library of THP-1 cells transduced with dCas9-KRAB, the reporter, and the hCRISPRi-v2 genome-wide sgRNA CRISPRi library (38). We infected this library with Mm (MOI 5), and 24 hpi used cell sorting to collect the highest ∼50% subpopulation (“reporter-high”) and the lowest ∼50% subpopulation (“reporter-low”) by reporter fluorescence (Fig. 1c). Genomic DNA was harvested and the guide-encoding region was amplified and deep-sequenced to identify enriched guides. Each gene was ranked for positive and negative enrichment using the MAGeCK analysis tool (39). In total, the screen detected 2035 significantly enriched genes (p<0.05), and 22 genes with false discovery rate (FDR) of <0.25, an FDR cutoff used in other exploratory CRISPR screens (40–42). The results of this screen represent a wide-range of potentially novel regulatory factors for IFN-β during infection.

Pathway- and gene-level analysis of the screen identified several known regulators of IFN-I, validating the screen (Fig. 1e; Fig. S1a). Top pathways in the reporter-low subpopulation included a variety of signaling pathways directly related to IFN-I signaling (Fig. 1d). In comparison, pathways from the reporter-high subpopulation were statistically weaker, and were dominated by mediator complex pathways, which mediates signals from transcription factors to RNA polymerase – potentially modulating transcription of interferon genes (Fig. S1a). Top reporter-low genes included the known IFN-β regulators, STING, STAT2, IRF9, and IFNAR1, suggesting the reporter was activated by both intrinsic and extrinsic, autocrine-paracrine, pathways (Fig. 1e and 1f). The top high-reporter genes included USP18, known to negatively regulate IFN-β, at rank 54 (8). Overall, the strong enrichment of known regulators of IFN-I signaling validates the methodology of the screen.

As the results of our screen included both known regulators of IFN-I and many unexpected genes, it suggests there are more regulators of IFN-I signaling than previously known. One of these unexpected genes was UFL1: The E3-like ligase of the UFMylation pathway.

### UFL1 suppresses transcription of IFN-β, IFIT1, and IL-6 during Mtb infection

To test these new putative regulators of IFN-β signaling, we chose eight genes enriched in the high-reporter subpopulation for testing via individual sgRNA-targeting, including the known IFN-β regulator, USP18, as a positive control. After infection and testing with the reporter, six of the targets did not give rise to a phenotype, although we did not test the efficacy of the CRISPRi-depletion and hence it’s not certain gene expression was perturbed (Fig. 2a). For the remaining two genes, UFL1 and NEDD8, CRISPRi-depletion resulted in increased reporter signal, although the increase for NEDD8 was not statistically significant. As the phenotype for NEDD8-deficiency was small and a role for NEDD8 in regulating IFN-β has been reported previously, we focused on UFL1 (43). Importantly, UFL1 had an FDR of 0.061 in our screen, which was within our preferred cut-off of 0.25 for significant enrichment.

**Fig. 2.**
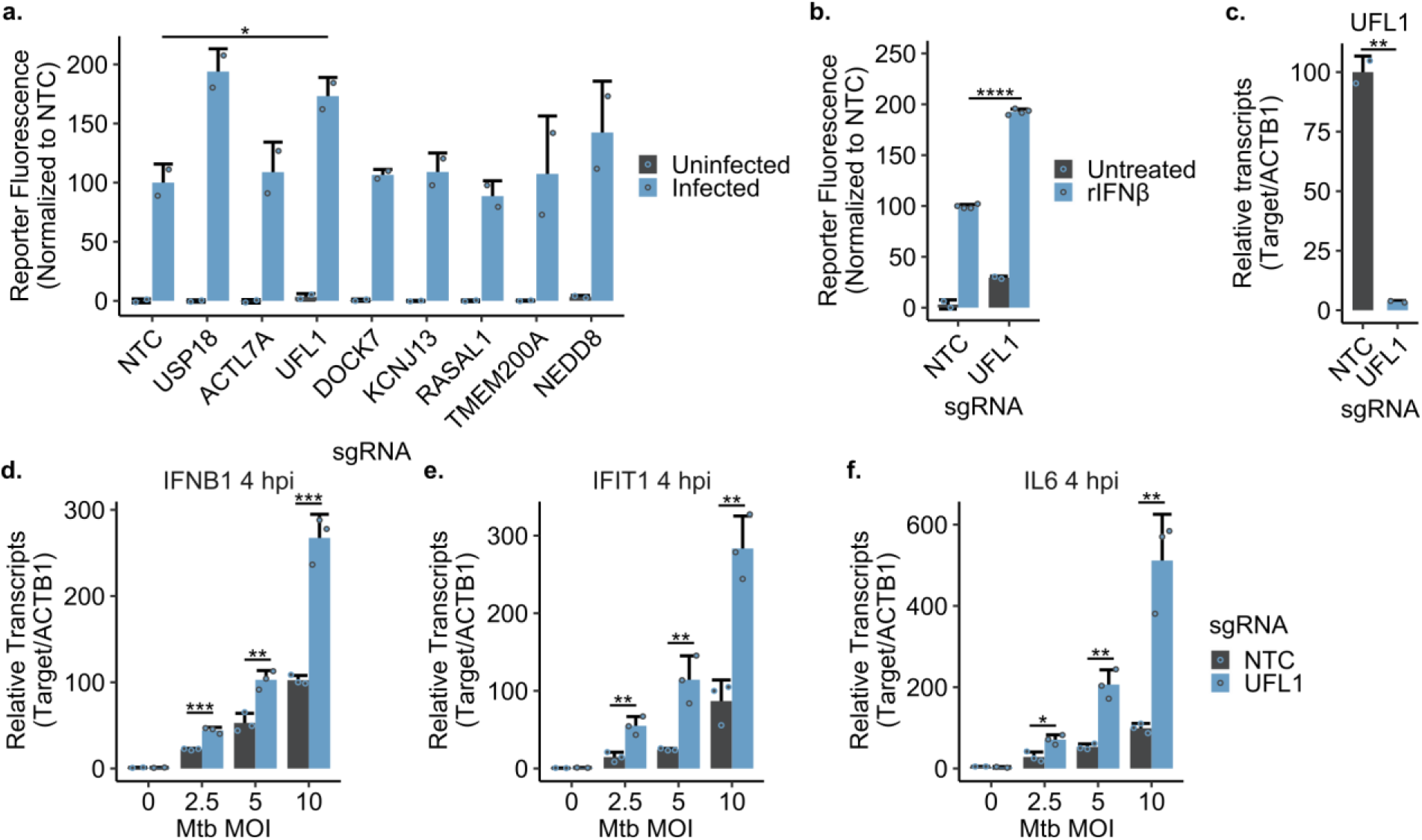
UFL1 suppresses transcription of IFN-β, IFIT1, and IL-6 during Mtb infection. (**a**). THP-1 macrophages with the indicated sgRNA were infected with WT *M. marinum*, MOI 5, or left untreated, and reporter expression was measured by flow cytometry at 24 hpi. NTC is Non-Targeting Control. (**b**). THP-1 with the indicated sgRNA were treated with 100 U/mL of human recombinant interferon beta, or left untreated. After 24 hours, reporter expression was measured by flow cytometry. (**c**). THP-1 with the indicated sgRNA were harvested for RNA and UFL1 transcripts were measured by RT-qPCR. (**d-g**). THP-1 with the indicated sgRNA were either left untreated or infected with WT *M. tuberculosis* (Erdman) at the indicated MOI for 4 hr, and transcripts for IFN-β, IFIT1, and IL-6 were measured by RT-qPCR. All results are representative of at least two independent biological experiments. Error bars represent SD with replicates n = 2 for a-c, n = 2 for d-f uninfected conditions, n = 3 for d-f infected conditions. *p<0.05, **p<0.01, ***p<0.001, ****p<0.0001 by unpaired t-test.

Because Mtb and Mm are known to stimulate production of IFN-I via the cGAS-STING- IRF3 pathway, we tested whether other pathways for IFN-I elicitation were altered by UFL1-deficiency. To test this, we treated UFL1-deficient and control cells with recombinant IFN-β, which signals through IFNAR1/2. Similar to the results with infection, we observed hyper-induction of the ISRE-IFN-β reporter in the UFL1-deficient cells relative to control cells (Fig. 2b). This suggests the effects of UFL1-deficiency on IFN-I signaling are not restricted to mycobacterial-stimulation of cGAS-STING signaling.

To support the conclusion that UFL1-deficiency was causing the enhanced reporter signal, we performed RT-qPCR, finding CRISPRi caused an approximately 97% depletion in UFL1 transcripts (Fig. 2c). This finding is consistent with UFL1-deficiency being the driver of the observed phenotypes, rather than off-target effects of CRISPRi (Fig. 3a).

**Fig. 3.**
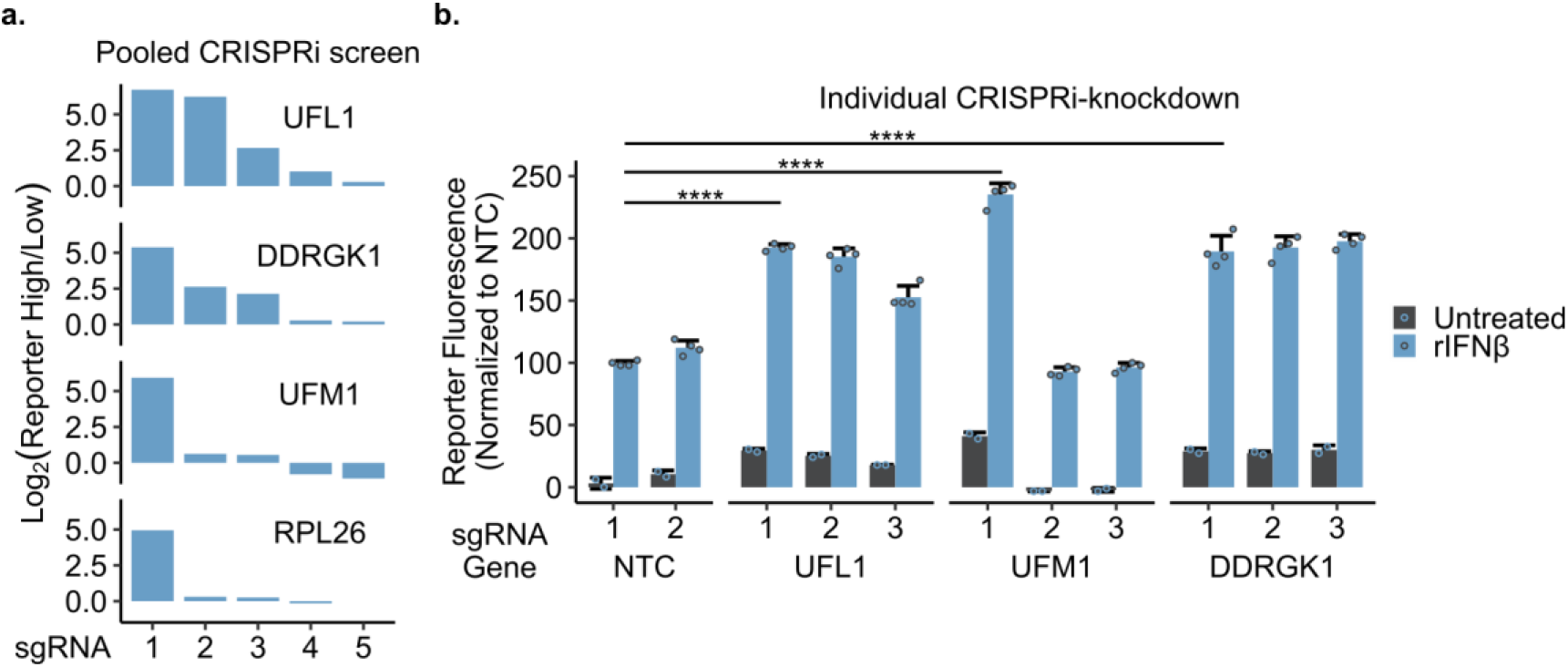
The UFMylation pathway inhibits immune signaling. (**a**) Log_2_-fold enrichment of sgRNA against the indicated gene from the pooled genome-wide CRISPRi screen. (**b**) THP-1 macrophages with indicated sgRNA were treated with 100 U/mL of human recombinant interferon beta or left untreated. At 24 hpi, reporter expression was measured by flow cytometry. NTC is Non-Targeting Control. Results are representative of at least two independent biological experiments. Error bars represent SD with replicates n = 2 for untreated and n = 4 for treated. *p<0.05, **p<0.01, ***p<0.001, ****p<0.0001 by unpaired t-test.

Next, we validated the reporter and screen findings with Mtb infection and RT-qPCR. In the uninfected condition, there was no difference in IFNB1 transcripts with UFL1- deficiency, however, during Mtb infection, UFL1-deficient macrophages had approximately 2.6-fold increase in IFNB1 transcripts relative to control cells at 4 hpi, MOI 10, and approximately 2-fold increases at MOI 5 and 2.5, 4 hpi (Fig. 2d). We hypothesized these cells would also hyper-induce ISGs after Mtb infection. Indeed, UFL1-deficiency increased transcripts of the ISG, IFIT1, by approximately 3-fold in RT-qPCR (4 hpi, MOI 10) (Fig. 2e). To determine whether the effect of UFL1-deficiency was specific to IFN-I, we measured transcripts for another pro-inflammatory cytokine, IL-6, and found that its transcripts were also increased approximately 5-fold with UFL1- deficiency at 4 hpi, MOI 10 (Fig. 2f). Overall, these findings support the conclusion that UFL1 suppresses IFN-I signaling during Mtb-macrophage infection, however, the findings for IL-6 suggest that the effects of UFL1-deficiency may be much broader than IFN-I signaling.

### The UFMylation pathway inhibits immune signaling

Next we addressed the question of whether UFL1 was acting independently of its known role in UFMylation to suppress IFN-I. UFMylation of target proteins requires the sequential activation and conjugation of ubiquitin-like UFM1 to target substrates via an E1 (UBA5), E2 (UFC1), and E3 (UFL1) in a pathway analogous to ubiquitin conjugation, with DDRGK1 recruiting UFL1 to the ER, where both localize (29, 30). Importantly, some guides against UFM1 and DDRGK1 were also strongly enriched in the reporter-high subpopulation – to a similar degree as guides against UFL1 – but the UFM1 and DDRGK1 genes did not reach high rankings due to low enrichment of other guides against these genes and our stringent analysis methods (Fig. 3a). We hypothesized that these guides had low enrichment due to poor knockdown efficacy, and that it is the UFMylation pathway that is important for IFN-β regulation rather than UFL1 functioning in a separate pathway. To test this, we depleted known UFMylation components. Attempts to deplete UBA5 and UFC1 yielded no viable cells, while depletion of DDRGK1 and UFM1 led to enhanced ISRE-IFN-β-reporter signal with IFN-β treatment, phenocopying UFL1-depletion, and indicating that UFM1-conjugation is important for IFN-β attenuation (Fig. 3b). We also considered what substrates of UFMylation may be relevant for suppressing immune signaling. One of the best understood substrates is the ribosomal protein RPL26 and, indeed, one guide against RPL26 was enriched in our screen, suggesting RPL26 could be a relevant substrate (Fig. 3b) (30, 44). However, attempts to deplete RPL26 yielded no viable cells, perhaps due to an essential role for RPL26 in ribosomal function. Additionally, as multiple guides against UFMylation components all resulted in enhanced reporter signal, it suggests the phenotypes are not due to off-target effects of CRISPRi (Fig. 3b). Taken together, the results suggest functional UFMylation – conjugation of UFM1 – is critical for IFN-β regulation, rather than UFL1 acting independently of UFMylation.

Due to the reported role for UFMylations in ER homeostasis, we hypothesized that ER-stress from UFMylation-deficiency may be the mediator of the enhanced immune response. It has been previously reported that UFMylation-deficient cells demonstrate ER-stress, and it has been reported that ER-stressors enhance the production of IFN-β with LPS. The enhanced production of IFN-β with ER-stressors was reported to be dependent on the ER-stress protein XBP-1, potentially via an XBP-1-binding site near IRF3 (30, 32, 45, 46). We hypothesized XBP-1 could be mediating the enhanced immune response observed in UFMylation-deficient cells. To test this, we sought to test UFL1-deficient cells versus cells deficient for both XBP1 and UFL1, however, this was inconclusive due to low efficacy with double CRISPRi targeting, and poor knockout efficiency for UFL1 with CRISPR-cutting.

### UFMylation suppresses inflammatory cytokine secretion during Mtb infection

As IL-6 transcripts were elevated during Mtb infection in UFMylation-deficient macrophages, we sought to further test the breadth of UFMylation’s effect on inflammatory signaling, and to confirm that the observed phenotypes extend to the protein level. To do this, we used a flow cytometry-based immunoassay to measure secreted cytokines in the supernatants from macrophages 24 hpi with Mtb. We observed approximately 2.5-fold and 40-fold more IFN-β and IL-6, respectively, from UFL1-deficient cells compared to control cells at MOI 5 (Fig. 4a and 4b). During infection, UFL1-deficient cells also secreted approximately 4-fold more TNF, and 2- to 4-fold more IFN-λ1, IFN-α2, IL-10, and IL-1β (MOI 5)(Fig. 4c-j). The results suggest an unexpectedly broad effect of UFMylation on inflammatory signaling.

**Fig. 4.**
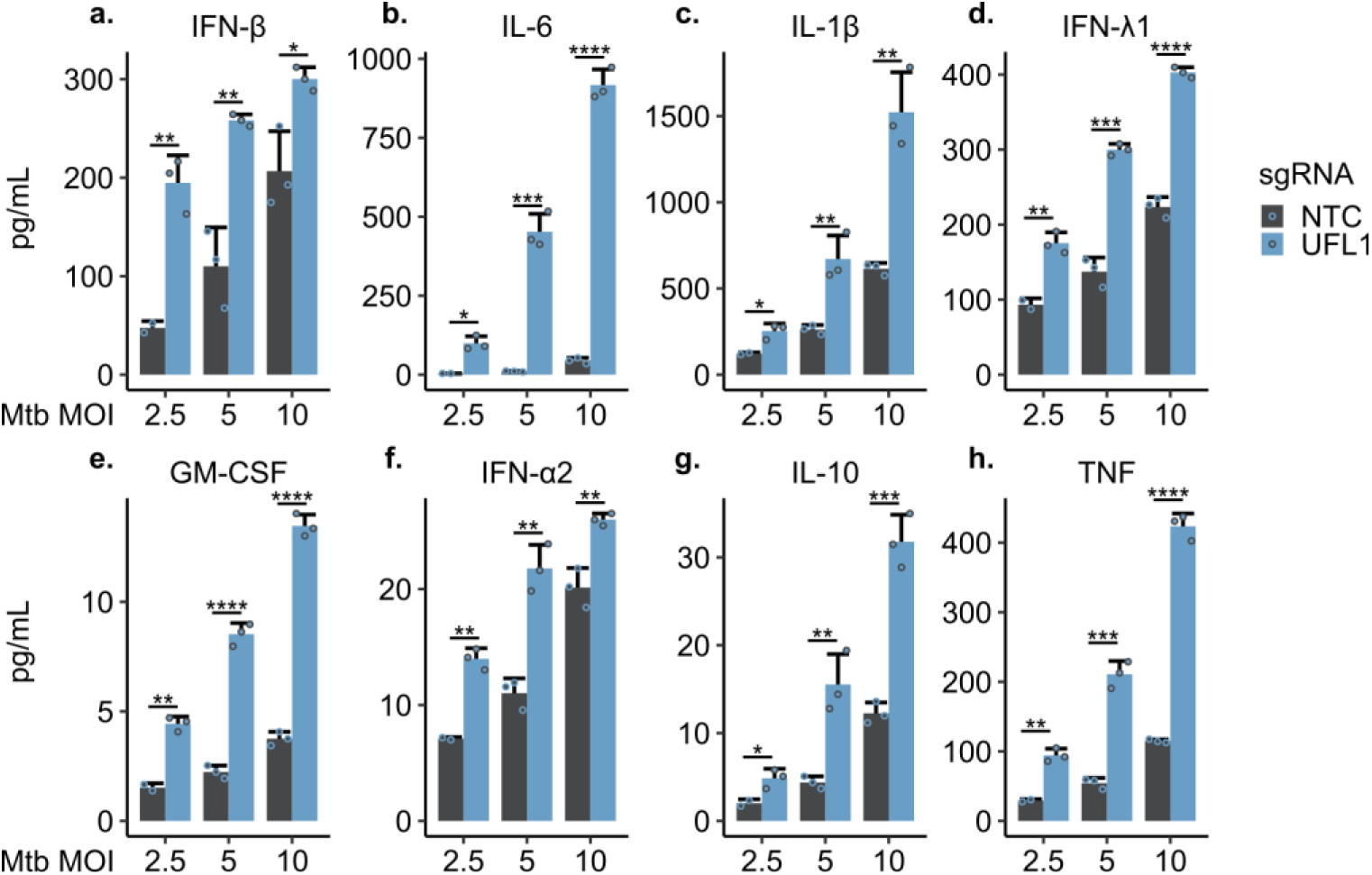
UFMylation suppresses cytokine secretion during Mtb infection. (**a-h**). THP- 1 macrophages with the indicated sgRNA were infected with WT *M. tuberculosis* (Erdman) at the indicated MOI for 24 hr. Supernatants were then harvested and protein levels were measured by cytometric bead array. NTC is Non-Targeting Control. Results are representative of at least two independent biological experiments. Error bars represent SD with replicates n = 3 for all except n = 2 for NTC MOI 2.5. *p<0.05, **p<0.01, ***p<0.001, ****p<0.0001 by unpaired t-test.

### Transcriptional profile of UFMylation-deficiency during Mtb infection

Our RT-qPCR and immunoassay results suggested that the effects of UFMylation on inflammatory signaling extend beyond inhibiting IFN-β production. To comprehensively determine how UFMylation alters the global macrophage response to Mtb infection, we performed RNA-seq on RNA from Mtb-infected UFL1-deficient and control cells at 4 hpi, MOI 10, and from uninfected cells.

Inspecting individual differentially-regulated genes showed that one of the most down- regulated genes was UFL1, indicating that the CRISPRi was effective (Fig 5a and 5c). Next we confirmed our prior results for IFN-β, finding an approximately 2.3-fold increase in transcripts for IFN-β from UFMylation-deficient cells compared to control cells during Mtb infection (Fig. 5e). Consistent with our prior findings, transcripts for IFIT1 and IL-6 were increased approximately 3-fold and 4.5-fold, respectively (Fig. 2e, 2f, and 5e). Also elevated were many other inflammatory genes, IL12B, MCP1, MCP2, and ACOD1 (3- to 9-fold increases), and a variety of ISGs, GBP1, IFIT2, IFIT3, and IFI44L (3- to 4-fold increases)(Fig. 5e). Notably, a number of genes were robustly induced by infection but did not appear strongly affected by UFL1-deficiency during infection, such as IL32, NINJ1, P47phox, LTB, and several genes associated with NF-κB signaling, such as NFKB2, NFKBIE, and RELB (Fig. 5a and 5e). Our findings from this gene-level analysis further indicate UFMylation has an unexpectedly broad, but not nonspecific, effect on the inflammatory response that extends beyond IFN-I signaling.

**Fig. 5.**
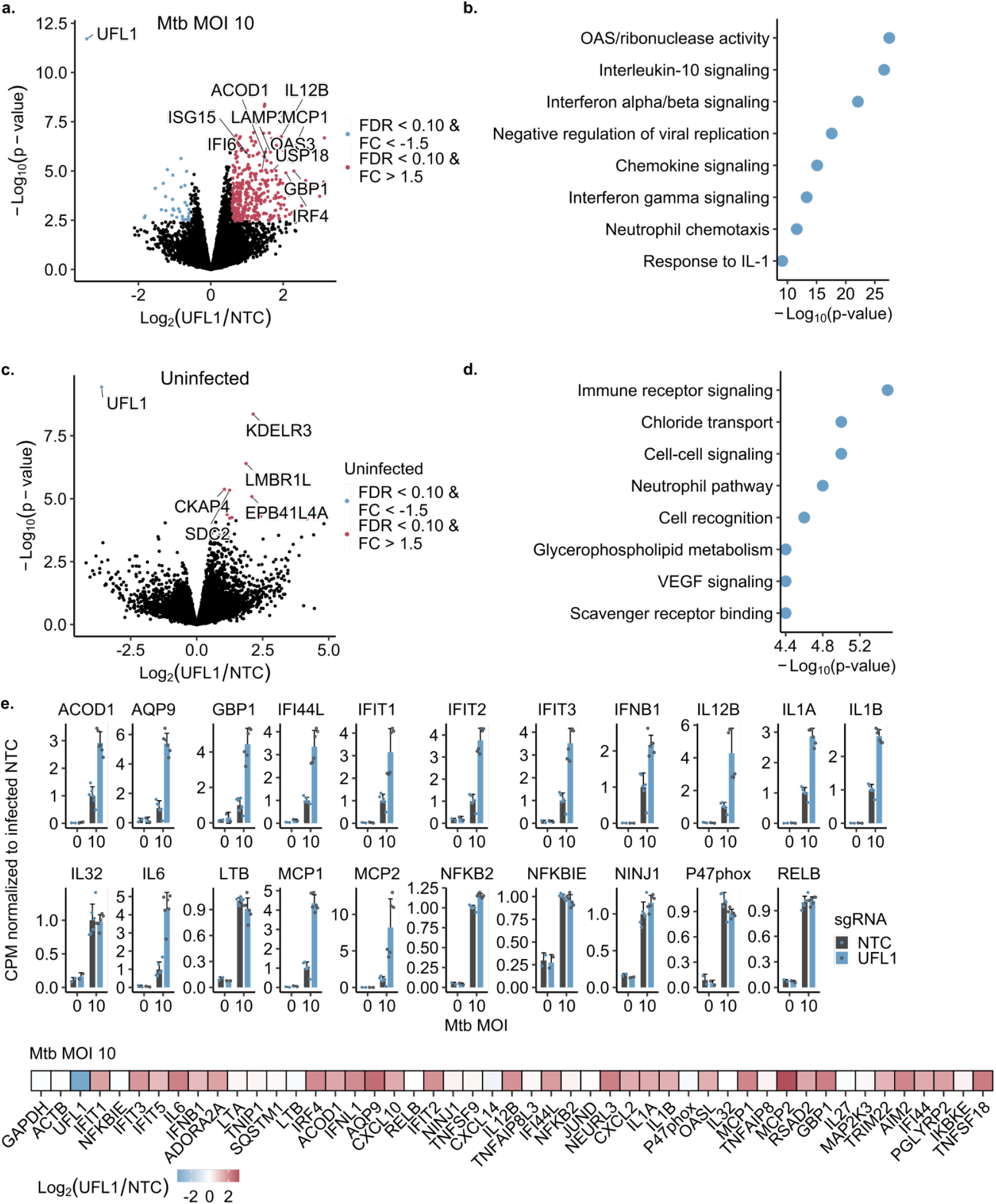
Transcriptional profile of UFMylation-deficiency during Mtb infection. (**a**). THP-1 macrophages with the indicated sgRNA were infected with WT *M. tuberculosis* (Erdman) at the indicated MOI. At 4 hpi, RNA was harvested and analyzed by RNA-seq. Results represented by volcano plot. NTC is Non-Targeting Control. FC is Fold Change. FDR is false discovery rate. Adjusted p-value (FDR) calculated by Benjamini-Hochberg procedure. (**b**). Pathway analysis of RNA-seq results, testing for pathway enrichment in genes differentially regulated by UFL1-deficiency (>1.5 fold change & p-value < 0.05) and induced by infection (>3-fold relative to non-infected cells). Some pathway names were shortened/simplified for brevity and some redundant pathways were removed/combined, keeping only the most statistically strong representative pathway. (**c**). Same as (a) but uninfected cells. (**d**). Pathway analysis of RNA-seq results from uninfected cells, testing for pathway enrichment in genes differentially regulated by UFL1-deficiency (>1.5 fold change & p-value < 0.05). (**e**). Gene-level results for RNA-seq. Top: Values as counts per million (CPM) normalized to the average of infected NTC per independent experiment. Error bars represent SD. Bottom: Heatmap showing differential regulation of genes induced by infection at least 3-fold. All results are pooled data from two independent biological experiments with replicates n = 6 for infected conditions, n = 2 for uninfected conditions.

Pathway analysis identified pathways enriched in genes that were most differentially upregulated with UFL1-deficiency and induced by Mtb infection. The top ranked pathways were 2’-5’-oligoadenylate synthetase activity, IL-10 signaling, interferon alpha/beta signaling, and chemokine signaling (Fig. 5b). The IL-10 set was attributed to upregulation of a broad list of inflammatory genes that included IL1A, IL6, IL12B, and TNF, while the interferon alpha/beta signaling set was attributed to IFNB1 along with several ISGs, such as IFITs/IFIs. The 2’-5’-oligoadenylate synthetase set was attributed to OAS1/2/3 genes, while chemokine signaling was attributed to upregulation of over a dozen chemokines, such as CXCL2 and CXCL10. Thus, pathway analysis supports our prior gene-level analysis of the broad effects on the inflammatory response by UFMylation.

Overall, transcriptional profiling suggests that UFMylation-deficiency caused an upregulation of a wide-variety of inflammatory genes and pathways, indicating UFMylation functions to broadly dampen inflammatory immune signaling during Mtb macrophage infection.

### Transcriptional profile of UFMylation-deficiency in the absence of infection

Transcriptional profiling of UFMylation-deficient cells versus control cells in the absence of infection revealed upregulation of many ISGs, including IFI30, IFI44L, and OAS1, although interferon-related pathways were not amongst the top enriched pathways. Pathway analysis revealed the most enriched pathway to be “immune response-regulating cell surface receptor signaling pathway,” attributed to enrichment of the genes CD24, FPR2, and CD40, which are involved in inflammation regulation (Fig. 5c and 5d). Overall, pathway analysis was much weaker statistically for uninfected cells, and hence major transcriptional effects caused by UFMylation-deficiency appear to primarily emerge with an additional stimulus like Mtb infection.

In conclusion, these findings suggest UFMylation functions to broadly dampen inflammatory immune signaling, with the most pronounced effects emerging during Mtb infection.

## DISCUSSION

While IFN-I appears to play a critical role in effective anti-viral immunity, evidence from human and mouse studies demonstrate that IFN-I can hinder control of Mtb infection (4, 5, 7–12, 15, 16). To determine genes that modulate IFN-I during mycobacterial infection, we performed a genome-wide CRISPRi screen in human THP-1 cells. The screen identified a number of known IFN-I regulators, such as STING and USP18, but also potentially novel regulators like UFL1, and revealed a role for UFMylation in suppressing IFN-β and inflammatory responses during Mtb infection (8, 24, 29).

Our findings contrast with Tao et al., which found siRNA-depletion of UFL1 in murine macrophages and human epithelial cells caused decreased production of IFN-β during viral infection – with UFL1 regulating STING independently of UFMylation (31). These differences may reflect differences in cell type, methodology (siRNA versus CRISPRi), or pathogen-specific virulence factors.

To address how UFMylation modulates immune responses, we consider one of UFMylation’s substantiated roles: ER homeostasis. One possible model for our findings is that ER-stress potentiates the IFN-I response elicited by Mtb, although many other models may be just as likely (30, 44, 47–50). UFMylation is reported to have a major role in resolving stalled ribosome-translocon complexes, with defects in UFMylation leading to ER-stress (30, 44, 45, 47, 50). Importantly, it has been reported that immune cells synthesize greater quantities of IFN-β in response to TLR ligands when treated with ER stressors (32). This was dependent on Xbp1, suggesting that Xbp1 may be mediating enhanced IFN-β in UFMylation-deficient cells. However, our experiments to test cells deficient for both XBP1 and UFL1 were inconclusive due to poor knockdown efficacy with double CRISPRi targeting, and poor knockout efficiency for UFL1 with CRISPR-cutting, leaving the question for future studies to resolve.

Because the findings reported here were made using the THP-1 cell line exclusively, it will be important to show that the regulatory effects of UFMylation extend to animal models of TB infection and ultimately human cells. Importantly, our results are consistent with work by Balce et al. which showed that UFMylation-deficiency also resulted in enhanced inflammatory signaling in response to IFN-γ treatment in murine BV-2 microglial cells, indicating that UFMylation-dependent regulation of immune responses is not specific to THP-1 cells (45). While UFMylation-deficiency is lethal in mammals, conditional, tissue-specific UFMylation-deficient mice have recently been generated that will allow for testing the role of UFMylation during Mtb infection *in vivo* (45, 51). As IFN-I signaling antagonizes Mtb control, a simple prediction is that UFMylation-deficiency would promote Mtb disease *in vivo* due to increased IFN-β. However, our findings indicate that UFMylation-deficiency has pleiotropic effects on macrophage immune signaling including increased production of several pro- inflammatory cytokines known to promote anti-Mtb immunity (e.g. TNF, IL-6, and IL-1) (52–54). Furthermore, even increased production of protective cytokines can have inverse effects on Mtb control *in vivo* – for example, excessive TNF causes harmful necroptosis and cachexia, high IL-6 is associated with increased tissue damage, and exuberant IL-1 signaling can lead to a harmful neutrophil cascade, demonstrating that their protective effects require precise regulation (55–62). Likewise, UFMylation- deficiency has also been reported to enhance responsiveness to IFN-γ, another pro- inflammatory cytokine critical for Mtb control (45, 52, 63, 64). Taken together, given the precise control of immune signaling required to control Mtb infection, it seems likely that UFMylation-deficiency would lead to exacerbated disease *in vivo*. Given the polyphenic nature of UFMylation in immune signaling, determining the exact mechanisms that may control Mtb growth and disease will be a formidable task. Finally, in human populations, variants in UFMylation components have been associated with severe health and developmental problems, particularly neurodegenerative diseases like microcephaly and epileptic encephalopathy (65). Although, we were unable to identify polymorphisms that would affect UFMylation genes in publicly-available association studies on Mtb susceptibility provided by the NHGRI-EBI GWAS Catalog (66). However, there is currently very limited information available on common human polymorphisms in UFMylation components, and it remains possible that advances in our understanding of polymorphisms in UFMylation components will reveal associations with Mtb pathogenesis.

As IFN-I are known to have antiviral properties, we also hypothesized that UFMylation- deficiency could be protective during viral infection. Indeed, loss of UFMylation inhibits hepatitis A virus (HAV) translation and HAV infection (67). The mechanism behind this observation is not fully elucidated, but repressed viral translation with loss of UFMylation is consistent with our findings of enhanced antiviral IFN-I with UFMylation-deficiency.

Next, we consider several speculative models to explain why UFMylation modulates IFN-I signaling. Our first model considers reports that viral infection causes ER-stress and that ER-stress enhances IFN-I production – hence ER-stress could function to detect and respond to viral infection (32, 68). In this system, normal ER function must be regulated by homeostatic mechanisms like UFMylation. If viral infection tips the scales past what these homeostatic systems can tolerate, ER-stress could enhance antiviral responses to counteract the infection. Consequently, UFMylation could fulfill an essential role in maintaining this antiviral surveillance system.

In another model, we consider that IFN-I have anti-neoplastic effects and UFMylation has several reported links to tumor surveillance and DNA-damage: Studies found that UFMylation promotes stability of the tumor suppressor p53, and histone UFMylation was found to be an important step for activation of the DNA damage response kinase ATM (69, 70). It’s then possible that UFMylation safeguards against tumorigenesis but, if lost, could potentiate IFN-I anti-tumor responses. While past studies found increased tumorigenesis in mice with UFMylation-deficiency, this was performed with tumor cell lines reportedly insensitive to IFN-β (69, 71). Therefore, our findings suggest a potential role for UFMylation in modulating IFN-I for tumor surveillance.

A third model to explain the connection between UFMylation and IFN-I signaling is that UFMylation could fulfill a guard role against viral activity: If viral activity inhibits UFMylation, that would serve as a warning to enhance antiviral responses. While UFMylation components were not decreased in HSV-1 and SARS-CoV-2 infection, it is not impossible that other viruses degrade UFMylation components, or that some viral factors inhibit UFMylation without degradation (72, 73). Alternatively, UFL1 could be sequestered from the ER due to viral activity – precluding UFMylation function at the ER – which, speculatively, could be achieved by virus-induced DNA damage. In the absence of infection, UFL1 was reported to localize to the ER (30). However, UFL1 also has a reported role in UFMylating nuclear proteins in response to DNA damage (69, 70). To fulfill this role, UFL1 must localize to the nucleus, potentially precluding UFL1’s ER functions. Numerous viruses are reported to cause DNA damage, and if this recruits UFL1 to the nucleus, this could mimic UFMylation-deficiency, enhancing IFN-I signaling (74).

Our work reveals a link between UFMylation and IFN-β during Mtb infection. Further investigation of UFMylation and IFN-I signaling during Mtb infection may determine the mechanism by which UFMylation regulates immune responses, and how UFMylation affects the outcome of Mtb infection. These studies will provide important insights into how IFN-I signaling impacts Mtb disease, and how we may be better able to treat Mtb by manipulating regulators of IFN-I signaling such as the UFMylation pathway.

## Supplementary Figures

**Fig. S1.**
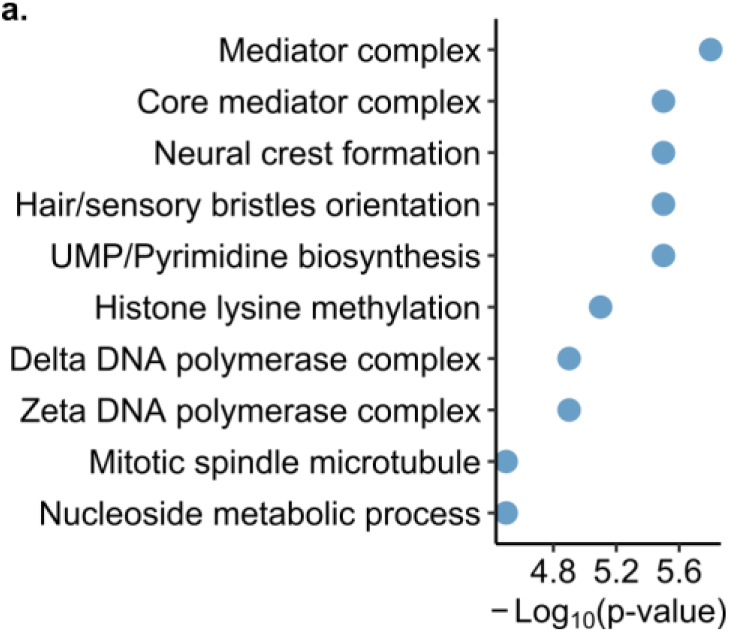
CRISPR interference screen for regulators of IFN-β during mycobacterial infection. (**a**) Gene set enrichment analysis of CRISPRi results, testing for enrichment in the top 250 genes for the reporter high population as ranked by Robust Rank Aggregation with MAGeCK. Some pathway names were shortened/simplified for brevity and some redundant pathways were removed/combined, keeping only the most statistically strong representative pathway.

## Methods

### RT-qPCR

Cells were harvested in TRIzol, RNA was isolated using PureLink RNA-Mini Kits (Invitrogen), treated with TURBO-DNase (ThermoFisher), and reverse-transcribed with SuperScript III cDNA Kit (ThermoFisher) with 500 ng of RNA. RT-qPCR was performed using SsoAdvanced SYBR Green-Supermix (Bio-Rad).

### RNA-seq

>150 ng of DNase-treated RNA, as described for RT-qPCR, was submitted to Novogene for rRNA depletion, library preparation, and sequencing with an Illumina NovaSeq X Plus (150 paired-end reads, for >20M reads per direction per sample). Mapping by Novogene. The edgeR and Limma R packages were used for normalization and statistical analysis with Voom.

### Mycobacterial culture

*M. tuberculosis* (Erdman) and *M. marinum* (M) were used. Mtb was grown to log phase in 7H9 liquid media (BD) with 10% Middlebrook OADC (Sigma), 0.5% glycerol, 0.05% Tween80 in roller bottles at 37°C. Mm was grown similarly except 0.20% Tween80 and 30°C.

### Macrophage infection

Macrophages were infected as previously described (75). Briefly, cultures were washed twice with PBS. Aggregates were removed by low-speed centrifugation (60xg, 5 min) and gentle sonication. Cells were overlaid with the bacterial suspension and ‘spinfected’ (1,000 xg, 10 min).

### Reporter assays

The reporter line was generated and used as previously described (34). Briefly, after 24 hours of treatment, reporter expression was measured in the PE channel with a BD LSR Fortessa. Fluorescence values were normalized to control cells by subtracting basal fluorescence of the untreated control cells, and expressing fluorescence as a percentage of the treated control cells.

### Lentiviral transductions

CRISPR guides derived from the CRISPRiv2 library (38). Oligonucleotides for sgRNAs were cloned into pLentiGuide-Puro (Addgene #52963) and checked by Sanger sequencing. 293T cells were lipofected with pLentiGuide-Puro, psPAX2, and pMD2.G using Lipofectamine3000 per manufacturer’s instructions. Target cells and lentivirus were spinfected (1000xg, 2 hr, 32°C) with 10 µg/ml protamine sulfate. Two days post-transduction, cells were selected with 4 µg/ml puromycin for ≥3 days.

### Cell culture

THP-1 were cultured in RPMI (Gibco) with 10% FBS and 5 mM L-glutamine. THP-1 were differentiated with 20 ng/mL of Phorbol-12-myristate-13-acetate (PMA) for 3 days and 1 day without prior to assays.

### CRISPRi screen

The THP-1 CRISPRi library was generated as previously described (34). Briefly, the THP-1 reporter cell line was transduced with the human genome-wide hCRISPRiv2 library containing 5 sgRNAs/gene. Cells were transduced and propagated at least 5×107 transduced cells, representing 500X coverage. The transduction efficiency was optimized for 20% to minimize multiple integrations per cell. Cells were infected with Mm as previously described. 24 hpi, cells were sorted using the BD Influx cell sorter and BD FACSaria Fusion cell sorter. After gating to remove debris and select GFP+/Reporter+ cells, cells were sorted into the highest 50% and lowest 50% of tdTomato-expressing cells. Genomic DNA was isolated using NucleoSpin Blood kits (Macherey-Nagel) and digested for 18 hours with SbfI-HF (NEB). The sgRNA cassette was isolated by gel electrophoresis with the NucleoSpin Gel and PCR Clean-up kit (Macherey-Nagel), and the DNA was PCR-amplified using primers with Illumina sequencing adapters. Indexed samples were sequenced via Illumina HiSeq-4000. Fastx-toolkit was used to generate the reverse complement of sequencing files where necessary. The screen was done for 3 independent biological replicates with a cumulative coverage of >500X. Results were analyzed using MAGeCK (39). Neg.score and alphamean from MAGeCK’s robust rank algorithm was used to quantify sgRNA enrichment with non-targeting sgRNAs as a control.

### Pathway analysis and volcano plots

The MAGeCKFlute package was used for pathway analysis with the hyper-geometric test and the GO, REACTOME, and BIOCARTA gene sets (76). For the CRISPRi screen, genes with a rank of ≤250 were used. For the RNA-seq pathway analysis, genes with ≥1.5-fold change and a p-value < 0.05 were used, along with a ≥3-fold increase relative to noninfected control cells for infected condition. Some pathway names were simplified for brevity and some redundant pathways were combined, keeping only the most statistically strong representative pathway.

### Cytokine analysis

Supernatants were filtered (0.22 um) twice. Cytokines were measured using BioLegend LegendPlex Plate (Hu Anti-Virus Response 13-plex) per manufacturer’s instructions.

